# Scientometric correlates of high-quality reference lists in ecological papers

**DOI:** 10.1101/2020.03.27.011106

**Authors:** Stefano Mammola, Diego Fontaneto, Alejandro Martínez, Filipe Chichorro

## Abstract

It is said that the quality of a scientific publication is as good as the science it cites, but the properties of high-quality reference lists have never been numerically quantified. We examined seven numerical characteristics of reference lists of 50,878 primary research articles published in 17 ecological journals between 1997 and 2017. Over this 20-years period, there have been significant changes in reference lists’ properties. On average, more recent ecological papers have longer reference lists, cite more high Impact Factor papers, and fewer non-journal publications. Furthermore, we show that highly cited papers across the ecology literature have longer reference lists, cite more recent and impactful papers, and account for more self-citations. Conversely, the proportion of ‘classic’ papers and non-journal publications cited, as well as the temporal range of the reference list, have no significant influence on articles’ citations. From this analysis, we distill a recipe for crafting impactful reference lists.

## Introduction

As young scientists moving our first steps in the world of academic publishing, we were instructed by our mentors and supervisors on the articles to read and cite in our publications. “Avoid self-citations”, “Include as many papers published in *Nature* and *Science* as possible”, “Don’t forget the classics”, and “Be timely! Cite recent papers” are all examples of such advices found in textbooks and blogs about scientific writing. Yet, to the best of our knowledge, intrinsic properties of high-quality reference lists have never been numerically quantified.

The success of a scientific publication varies owing to a range of factors, often acting synergistically in driving its impact. Apart from the scientific content of the article itself, which ideally should be the only predictor of its impact, factors that correlate to the number of citations that an article accumulates over time include its accessibility ^1,2^, the stylistic characteristics of its title ^3–5^ and abstract ^6^, the number of authors ^7^, and its availability as a preprint ^8^. Furthermore, it is understood that the quality of a scientific publication should be related to the quality of the science it cites, but quantitative evidence for this remains sparse ^7,9–11^.

From a theoretical point of view, a reference list of high quality should be a balanced and comprehensive selection of up-to-date references, capable of providing a snapshot of the intellectual ancestry supporting the novel findings presented in a given article ^12^. This is achieved by conducting a systematic retrospective search to select all papers with content that is strongly related to that of the article, to be read and potentially cited if deemed relevant. The most throughout and recent attempt to evaluate the quality and properties of a journal article reference list was made by Evans ^9^. Using a database of >30 million journal articles from 1945 to 2006 Evans showed that, over time, there has been a general trend to referencing more recent articles, channelling citations toward fewer journals and articles, and shortening the length of the reference list. Evans predicted that this way of citing papers “[…] *may accelerate consensus and narrow the range of findings and ideas built upon*”, an observation that generated subsequent debate ^13–15^. For example, in a heated reply to Evan’s report, Von Bartheld et al. ^13^ argued that this claim was speculative because “*[…] citation indices do not distinguish the purposes of citations*”. In their view, one should consider the ultimate purpose of each individual citation and the motivation of authors when they decided which papers to cite.

Yet, it is challenging to disentangle all factors driving an author choice of citing one or another reference ^11,16^, especially when dealing with large bibliometric databases such as the one used by Evans ^9^ to drawn his conclusions. In spite of the attempts made, the question remains as to how to objectively evaluate the quality and properties of a reference list. To address this gap, we extracted from Web of Science (Clarivate Analytics) all primary research journal articles published in low- to high-rank international journals in ecology in the last 20 years, and generated unique descriptors of their reference lists. We restricted our analysis to articles published in international journals in “Ecology” because, by focusing on a single discipline, it was possible to minimize the number of confounding factors. Moreover, this choice allowed us to incorporate in the analyses a unique descriptor of the reference list based on an analysis published in 2018 on seminal papers in ecology ^17^ (see “Seminality index” in Table 1).

**Table 1.** Proxy variables used to characterized the reference list of the papers.

| Variable | Description | Construction | Type |
| --- | --- | --- | --- |
| Length | Total number of reference items cited in the paper reference list. | Sum of reference items cited (variable "NR" in WoS database). | Real value |
| non-ISI papers (non-journal publication cited) | Number of non-ISI items cited in the reference list, such as books, theses, websites, and grey literature. | Number of non-journal items cited divided by Length. | Proportion |
| Self-citations | Number of self-citations in the reference list. | Number of self-citations divided by Length.<br>Note that only first author self-citations are counted, namely those in which any of the authors of the paper appear as first author in items of the reference list. | Proportion |
| Temporal span | Temporal span of the reference list. | Year of most recent reference item cited – year of oldest item cited. | Real value |
| Immediacy | Number of recent reference items cited. | Number of papers published in the previous three years divided by Length. | Proportion |
| Seminality | Number of seminal ecological papers cited | Number of cited items in the list "*100 articles every ecologist should read*"<sup>17</sup> divided by Length. | Proportion |
| Total IF | Sum of the IF values of all the reference cited in the paper. | Total IF is calculated using the JIF of each reference at the year of publication, based on annual JCRs. To calculate the proportion, Total IF is divided by the number of reference with JIF (Length – number of non-ISI items). | Proportion |

We structured this research under two working hypotheses. First, if the quality of a scientific paper is connected to the reference it cites, we predict that, on average, articles characterized by a good reference list should accumulate more citations over time, where the goodness of a reference list is approximated via a combination of different indexes (Table 1). Second, we hypothesize that thanks to modern searching tools such as large online databases, bibliographic portals, and hyperlinks, the behavior through which scientists craft their reference lists should have change in the Internet era ^15,18^. Thus, we predict that this change should be reflected by variations through time in the proprieties of articles’ reference lists.

## Results

### Reference list characteristics in ecology

After excluding non-primary research articles and omitting incomplete WoS records, we ended up with 50,878 unique papers distributed across the 17 journals that covered the time span from 1997 to 2017. The median size of the reference list in ecological journals is 54 cited items (range= 1–403) (Fig. 1a). Cited references cover a median temporal span of 45 years (0–922) (Fig. 1b). The mean proportion of recent papers in the reference lists is 0.21 (0–1); the proportion of non-ISI articles is 0.12 (0–0.8), whereas the average impact factor of the papers cited in references lists is 4.9 (0–29.5) (Fig. 1). The mean proportion of self-citations is 0.07 (Fig. 1f) and the proportion of cited seminal papers is 0.006 (0–0.33) (Fig. 1g).

**Figure 1.**
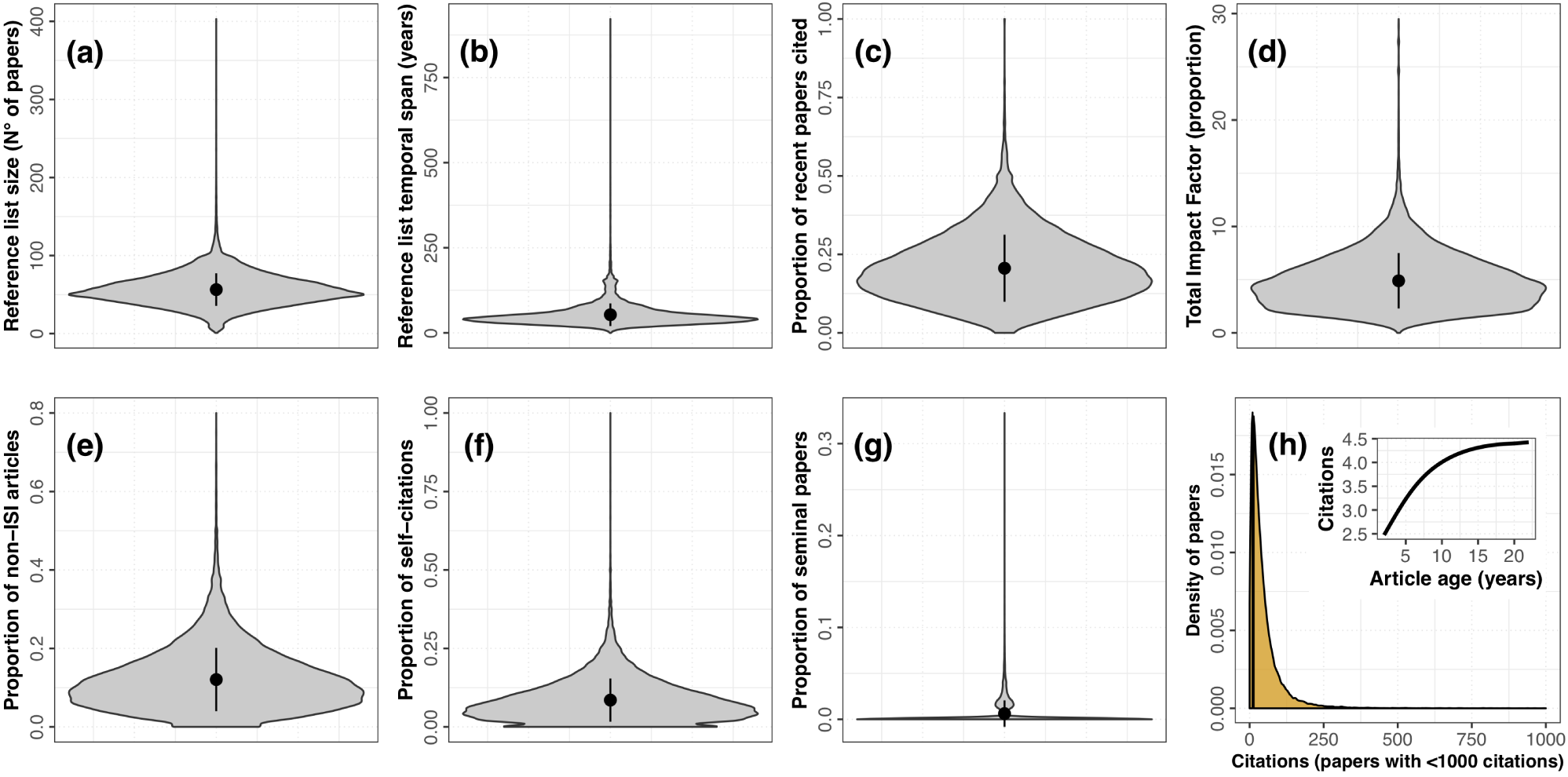
Main numerical features of reference list of ecological journals. **a–g**) Violin plots showing the distribution of the seven numerical properties of reference lists considered in this study. For each graph, black dot and vertical bar is mean ± s.d. **h**) Distribution of citations among the articles considered in this study. Inset show the predicted relationship between citations and articles age, based on the prediction of a generalized additive model.

We predicted the expected curve of citations over article age with a Poisson generalized additive model (GAM). We observed a significant parabolic trend in the number of citations over time (F= 2724.8; p< 0.001), with number of citations reaching a plateau of ∼4 after 10 years from publication (Fig. 1h, inset).

### Relationship between reference list features and article impact

To normalized the number of citations for each article by its age, we expressed citations as the Pearson residuals from the regression curve shown in Fig. 1h (inset). We modeled residuals of citations as a function of the different features of the reference list, using a linear mixed effects model with journal identity and publication year as random factors.

We observed a positive and significant relationship between citations of a paper and the number of cited references (Estimated β ± s.e. 3.11±0.12 p< 0.001), with articles with longer reference lists accumulating more citations over time (Fig. 2a). The number of citations also significantly increased with an increase in the proportion of self-citations (Estimated β ± s.e.: 3.45±0.34, p< 0.001; Fig. 2b) and reference list total Impact Factor (IF) (Estimated β ± s.e.: 0.99±0.12, p< 0.001; Fig. 2d). Furthermore, we found a positive relationship between citations and immediacy of the reference list, namely articles citing a greater proportion of recent papers accumulated more citations over time (Estimated β ± s.e.: 11.28±0.39, p< 0.001; Fig. 2c). Proportion of non-ISI journal article referenced, total temporal span of the reference list, and proportion of cited seminal papers had no significant effect on citations (non-ISI Estimated β ± s.e.: –0.22±0.39, p= 0.554; Temporal span: –0.13±0.35, p= 0.164; Seminality: 0.46±0.644, p= 0.470).

**Figure 2.**
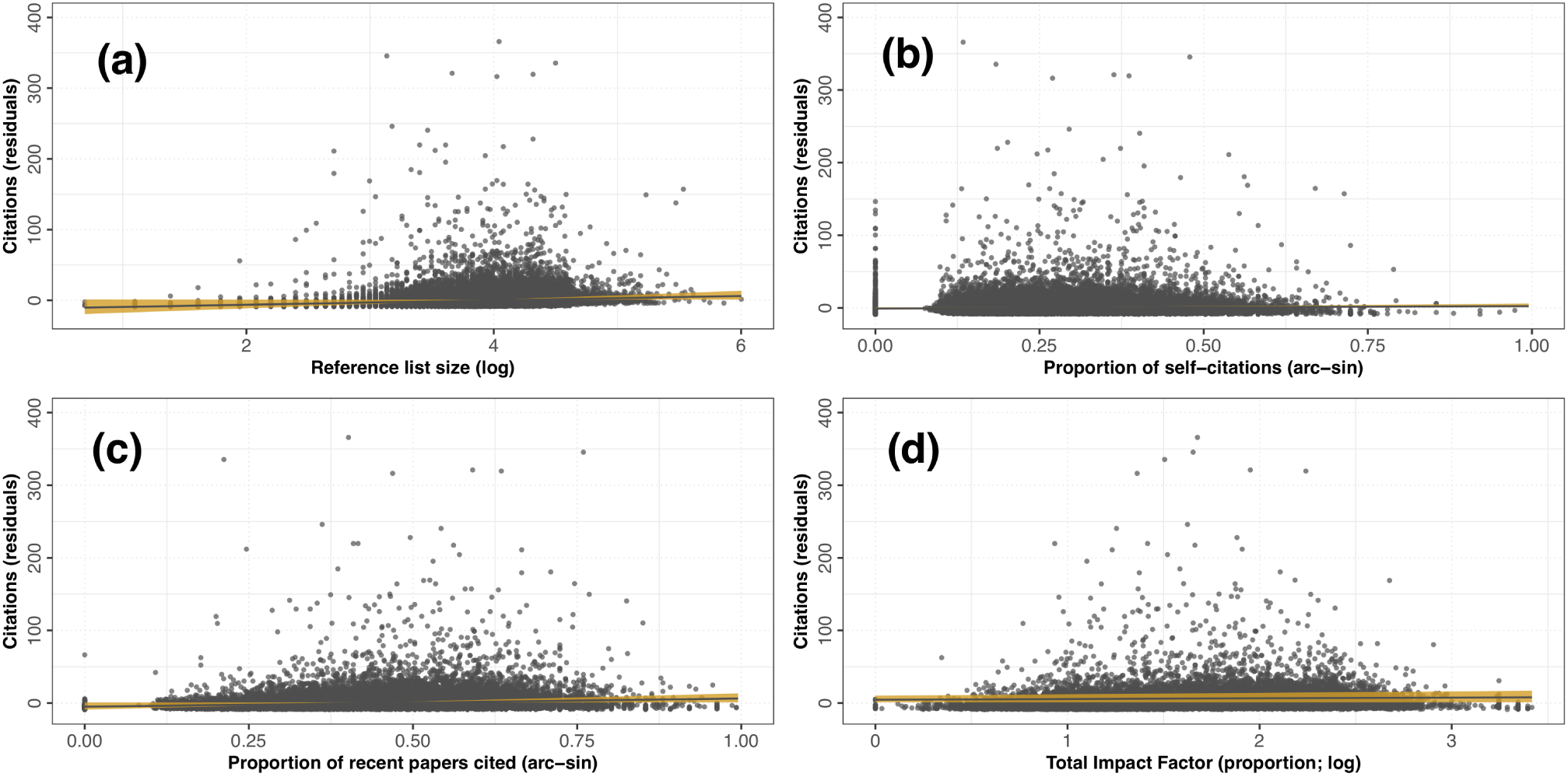
Relationships between articles citation and reference lists numerical properties. Predicted relationships (filled lines) and 95% confidence intervals (orange surfaces) between the residuals of citation over articles’ age and **a**) length of the reference list, **b**) proportion of self-citation, **c**) proportion of recent papers cited, and **d**) total impact factor of the reference list, according to the Linear mixed models analysis. Variables are transformed to homogenize their distribution. Only fixed effects are shown.

### Temporal variations in reference list features

Over the 20-years period considered (1997–2017), the total IF of the reference list steadily and significantly increased. The average (±s.d.) IF of articles cited in the reference list was 2.35±1.83 in 1997, and 6.19±2.23 in 2017 (Fig. 3c). Yet, it is worth noting that over the 20-years period considered the overall IF of scientific journals also significantly increased, a feature that may have inflated this trend ^19^. In parallel, the proportion of non-journal articles referenced significantly decreased over time. In 1997, on average, non-journal articles accounted for 14% of the reference list, while this value dropped to 8% in 2017 (Fig. 3b). We also observed that the number of cited items in the reference list steadily increased from an average of 45.3±20.5 in 1997 to 68.2±25.5 in 2017 (Fig. 3a). We observed stabler trends for the temporal span of the reference list (Fig. 3d), proportion of self citations (Fig. 3e), recent papers (Fig. 3f), and seminal papers (Fig. 3g). Models estimated parameters are in Fig. 3.

**Figure 3.**
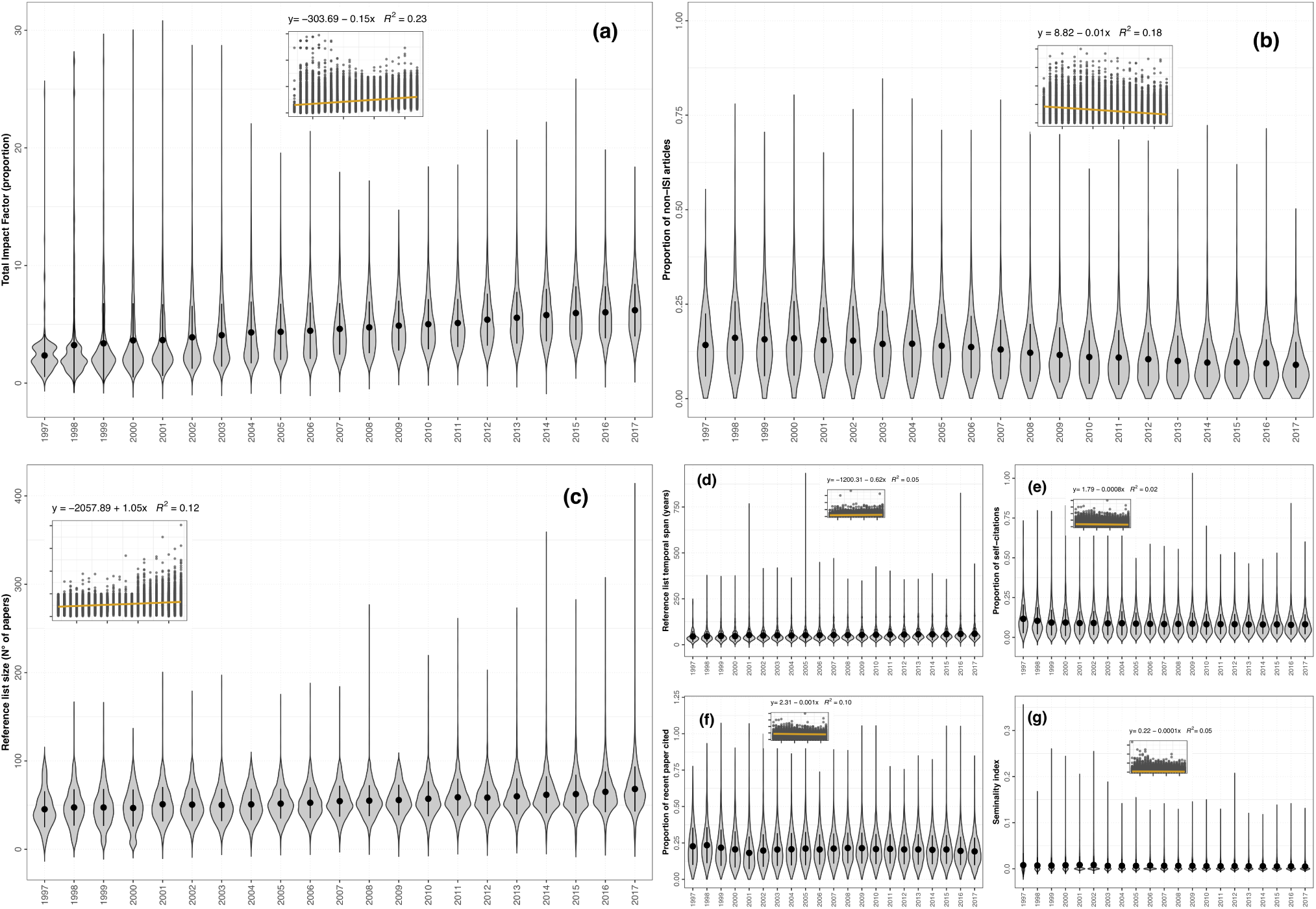
Variations in reference lists numerical properties between 1997 and 2017. a–g) Violin plots showing the annual variations in the seven numerical features of reference lists. Insets show the predicted relationships (filled line) and 95% confidence intervals (orange surfaces) based on linear mixed models. Larger graph (a–c) illustrate non-flat temporal trends. Only fixed effects are shown.

## Discussion

We showed that, on average, papers with longer reference lists are more cited across the ecological literature than papers with shorter reference lists, a result that parallels findings of previous studies ^7,9^. One explanation is that longer reference lists may make papers more visible in online searches. Also, it was hypothesized that papers with longer reference lists may address a greater diversity of ideas and topics ^7^, thus containing more citable information. Furthermore, a longer reference list may attract *tit-for-tat* citations, that is, the tendency of cited authors to cite the papers that cited them ^20^. It is interesting to emphasize that this result directly questions the practice of most journals to set a maximum in the number of citable references per manuscript. Since most journals are switching to online-only publishing systems where space limitation is not an issue, this limitation seems unjustified.

We also found that papers citing a greater proportion of recent articles and high-IF articles are, on average, more cited. Citing recent references generally implies that scientists are working on ‘hot’, timely eco-evolutionary topics. The latter frequently end up published in journals with greater impact factor, which on average attract a greater share of citations. A complementary explanation for this result may be searched for in the recent changes in academic publishing. It was pointed out that, since the volume of available scientific information in the Internet era is growing exponentially ^18,21^, scientists are not anymore able to keep pace with relevant papers published every year about any given scientific topic. As a result, they often end up reading almost exclusively the latest ‘hot’ papers ^17,18^ while avoiding older literature ^9^.

Furthermore, we found that papers including a greater proportion of self-citations are more highly cited. Given that excessive self-citations are usually despised and discouraged, this results may come at a surprise. On the one hand, it is true that self-citations are sometimes unjustified, used by authors as a way to increase their scientific visibility and to boost their own citation metrics ^10^. An irrelevant self-citation breaking the flow of a paragraph, such as this one^22^, is an instructive example of this behavior. On the other hand, self-citations are an integrant part of scientific progress, as they usually reflect the cumulative nature of individual research ^23^. Indeed, 88% of the papers in our dataset included at least one self-citation. This may ultimately lead to accumulate more citations, because papers that are part of a bigger research line are often more visible and citable.

According to our analyses, other features of the reference list have not significant effect on citations. Probably, the least intuitive result is a lack of relationship between the number of cited seminal papers and the number of citations. The list of seminal papers was generated using the results of a recent expert-based opinion paper, providing a list of the 100 “must-read” articles in ecology ^17^. A manuscript citing any of those classical papers should focus, on average, on broader and long-debate topics in ecology, and therefore it is expected to receive more citations. But this is not the case. If one assumes that the number of citations for a paper is an index of its importance for the field, such a result may question the “must-read” value of some of the articles included in Courchamp & Bradshaw ^17^ compilation. However, most of these seminal papers are relatively old and they thus have inspired more recent studies, which may be cited instead of the original ones.

### Change in reference lists structures over time

We observed significant changes in the structure of articles’ reference lists from 1997 to 2017. We argue here that most of these changes are directly related to a shift in the academic publishing behaviors of the Internet era ^24^ from browsing paper in print to searching online through the use of hyperlinks ^9,15^. While the volume of available scientific information has grown exponentially ^18^, retrieving relevant bibliography has become simpler and quicker thanks to online searching tools ^15^. This seemingly explain why, on average, the length of reference lists across ecological journals has steadily increased.

The last two decades have also seen an exponential rise in the use of journals metrics, especially the impact factor ^19^, and the consequent desire of authors to publish in high-ranking journals and cite papers published therein. This may explain why we observed a significant increase in the total impact factor of reference lists over time. Concomitantly, there has been a reduction in the number of non-ISI publications cited in reference lists. In general, both these features are a direct product of the changes in academic publishing behaviors of the “publish or perish” era. More and more authors are now exploring new ways to maximize the impact of their publications ^25,26^. Citing papers with higher impact factor and a lower proportion non-journal articles may be perceived as an effective way to achieve such goal.

## Concluding remarks

While we are writing, identifying and citing the most relevant articles that provide the scientific foundation for our research questions is not trivial. Time is against us: most researchers are overloaded by academic duties and have busy schedules, preventing to read classic papers and keep up with the latest advances in the main and nearby fields of research. Memory failures, perhaps increased by the haste of finishing the manuscript in time, do not help either. Accordingly, reference lists are almost inevitably characterized by faulty citations, including incorrect references, quotation errors, and omitted relevant papers ^16^. In a more cynical reasoning, May ^12^ even argued that omissions of relevant papers might be due to the simple fact that *“[…] the author selects citations to serve his scientific, political, and personal goals and not to describe his intellectual ancestry*.”

But once we accept that making the perfect reference list is not possible, three heuristic rules will help us getting close to it:

1. Size matters. Not only in terms of reference list, but also in the number of characters ^27,28^. Investing extra resources into reading others research it improves the scientific basis of the study while building argumentation links with relevant manuscripts, making the paper more visible and useful to peers.
2. Hotness. During the last twenty years we have seen the advent of the Internet and changes in the way information is found, read, and spread. Keep track of impactful latest research, even exploiting novel tools such as social media ^29^ and blogs ^30^, is a crucial premise to produce highly citable science.
3. Narcisism. Not only self-citations directly increase the citations of past work, but they have been shown to improve chances of being cited by others ^10^. Furthermore, the probability of self-citation increases with professional maturity in a given field of study, showing that that is a direct consequence of the cumulative nature of individual research ^23^.

## Methods

### Criteria for articles inclusion

We extracted from WoS all primary research articles published in the ecological journals between 1997 and to 2017 (Table 2). The year 1997 was chosen because approximately around this date the use of impact factor (IF) started to grow exponentially ^19^. We selected only those ecological journals covering more than 75% of the 20-years period considered, thus allowing to explore temporal trends with confidence. For example, *Nature Ecology and Evolution* (2016–ongoing) was excluded as it covered only 10% of this temporal interval. We selected exclusively primary research articles because review and opinion papers, methodological papers, corrections, and editorials may have atypical reference lists.

**Table 2.** Journal selected for the analysis.

| **Journal name** | **Initial year** | **Temporal span**<br>**selected** | **Totale N° of**<br>**articles** | **N° of primary research**<br>**articles** |
| --- | --- | --- | --- | --- |
| Acta Oecologica | 1983 | 1997–2017 | 1,571 | 1,408 |
| American Naturalist | 1867 | 1997–2017 | 3,417 | 2,852 |
| Austral Ecology | 2000 | 2000–2017 | 1,659 | 1,434 |
| Ecography | 1978 | 1997–2017 | 2,051 | 1,743 |
| Ecological<br>Applications | 1991 | 1997–2017 | 3,641 | 3,051 |
| Ecology | 1920 | 1997–2017 | 6,584 | 5,505 |
| Ecology Letters | 1998 | 1998–2017 | 2,636 | 2,098 |
| Functional Ecology | 1987 | 1997–2017 | 2,889 | 2,326 |
| Global Change<br>Biology | 1995 | 1997–2017 | 4,573 | 3,937 |
| Global Ecology and<br>Biogeography | 1993 | 1997–2017 | 1,570 | 1,377 |
| Journal of Animal<br>Ecology | 1932 | 1997–2017 | 2,639 | 2,250 |
| Journal of Applied<br>Ecology | 1964 | 1997–2017 | 2,993 | 2,407 |
| Journal of<br>Biogeography | 1974 | 1997–2017 | 3,541 | 2,852 |
| Journal of Ecology | 1913 | 1997–2017 | 2,603 | 2,170 |
| Molecular Ecology | 1992 | 1997–2017 | 7,853 | 6,209 |
| Oecologia | 1968 | 1997–2017 | 6,417 | 5,446 |
| Oikos | 1949 | 1997–2017 | 4,687 | 3,812 |

We generated seven descriptors of reference lists properties, and used these as variables in subsequent analyses. A description of each variable and the rationale for its construction are in Table 1. Note that most of the reference list descriptors are expressed as proportions, in order to normalize variables to the number of papers cited in the reference list ^31^.

### Relationship between citations and reference list characteristics

We conducted all analyses in R ^32^. To test our first working hypotheses, we conducted regression-type analyses following the general protocol by Zuur & Ieno ^33^. We initially explored our dataset following a standard protocol for data exploration ^34^, whereby we: i) checked for outliers in the dependent and independent variables; ii) explored the homogeneity of variables distribution; and iii) explored collinearity among covariates based on pairwise Pearson correlations—threshold for collinearity set at |*r*|> 0.7 ^35^.

As a result of data exploration, we removed three outliers from articles citations, corresponding to three papers cited over 6,500 times in WoS. We homogenized the distribution of our explanatory variables by log-transforming reference list size and temporal span, and square-root arcsin transforming all proportional variables. We also observed that over 40% of the articles in our dataset were never cited (Fig. 1a), but since these represent “true zeros” ^36^ we didn’t apply zero-inflated models to infer citation patterns over time ^37^. No collinearity was detected among the seven explanatory variables—all |*r*|< 0.7.

We used a Poisson generalized additive model (GAM) to predict the expected pattern of citations over article age, and expressed the number of citations as the Pearson residuals from the curve (Fig. 1a). To test which reference list features correlate with residuals in number of citations, we generated a linear mixed effects model (LMM) by including journal identity and publication year as random terms to account for data non-independence. We fitted LMM with the R package “nlme” ^38^, and validated models using residuals and fitted values ^33^.

### Change in reference list characteristics over time

We used LMMs to predict annual variations in reference list characteristics over time, including journal identity as a random factor. Seven LMMs were constructed, one for each variable described in Table 1. In these case, as the seven variables were included as dependent variables, we didn’t log-and square-root arcsin transformed variables.

## Acknowledgements

Thanks to Pedro Cardoso for a useful discussion.

## Author contributions

SM conceived the study. SM and FC designed methodology. FC mined data from WoS. SM and FC developed code for data processing. SM performed the analyses, with suggestions by FC, DF, and AM. SM wrote the first draft. All authors contributed to the writing of the manuscript through comments and additions.

## Data availability

All data used to generate this study can be freely downloaded from Web Of Science. The cleaned database and R script to generate the analysis will be deposited in a public repository upon acceptance of the peer-review version of this paper.

